# Biochemical characterization of the methylmercaptopropionate:cob(I)alamin methyltransferase from *Methanosarcina acetivorans*

**DOI:** 10.1101/548461

**Authors:** He Fu, Michelle N. Goettge, William W. Metcalf

**Author notes:** Corresponding author: Department of Microbiology, University of Illinois at Urbana-Champaign, B103 C&LSL, 601 South Goodwin Avenue, Urbana, IL, 61801. [a Current address: Department of Marine Sciences, University of Georgia, Athens, GA 30602.] [b Current address: Manus Bio 1030 Massachusetts Ave, Suite 300, Cambridge, MA, 02138].

## Abstract

Methanogenesis from methylated substrates is initiated by substrate specific methyltransferases that generate the central metabolic intermediate methyl-coenzyme M. This reaction involves a methyl-corrinoid protein intermediate and one or two cognate methyltransferases. Based on genetic data, the *Methanosarcina acetivorans* MtpC (corrinoid protein) and MtpA (methyltransferase) proteins were suggested to catalyze the methylmercaptopropionate(MMPA):Coenzyme M (CoM) methyl transfer reaction without a second methyltransferase. To test this, MtpA was purified after overexpression in its native host and characterized biochemically. MtpA catalyzes a robust methyl transfer reaction using free methylcob(III)alamin as the donor and mercaptopropionate (MPA) as the acceptor, with *k*_cat_ of 0.315 s^-1^ and apparent *K*_m_ for MPA of 12 μM. CoM did not serve as a methyl acceptor, thus a second, unidentified methyltransferase is required to catalyze the full MMPA:CoM methyl transfer reaction. The physiologically relevant methylation of cob(I)alamin with MMPA, which is thermodynamically unfavorable, could also be demonstrated, but only at high substrate concentrations. Methylation of cob(I)alamin with methanol, dimethylsulfide, dimethylamine and methyl-CoM was not observed, even at high substrate concentrations. Although the corrinoid protein MtpC was poorly expressed alone, a stable MtpA/MtpC complex was obtained when both proteins were co-expressed. Biochemical characterization of this complex was not feasible because the corrinoid cofactor of this complex was in the inactive Co(II) state and could not be reactivated by incubation with strong reductants. The MtsF protein, comprised of both corrinoid and methyltransferase domains, co-purifies with the MtpA/MtpC, suggesting that it may be involved in MMPA metabolism.

**IMPORTANCE:** MMPA is an environmentally significant molecule produced by degradation of the abundant marine metabolite dimethylsulfoniopropionate, which plays a significant role in the biogeochemical cycles of both carbon and sulfur, with ramifications for ecosystem productivity and climate homeostasis. Detailed knowledge of the mechanisms for MMPA production and consumption is key to understanding steady state levels of this compound in the biosphere. Unfortunately, the biochemistry required for MMPA catabolism under anoxic conditions is poorly characterized. The data reported here validate the suggestion that the MtpA protein catalyzes the first step in methanogenic catabolism of MMPA. However, the enzyme does not catalyze a proposed second step required to produce the key intermediate methyl-CoM. Therefore, additional enzymes required for methanogenic MMPA catabolism await discovery.

## INTRODUCTION

*Methanosarcina* species comprise a versatile group of methane-producing archaea capable of growth on a variety of methylated one-carbon (C_1_) compounds, including methanol, methylamines, and methylsulfides. Catabolism of these C_1_ substrates is initiated by transfer of the methyl moiety to coenzyme M (CoM, mercaptoethanesulfonate), generating methyl-CoM. This central metabolic intermediate is then disproportionated to produce carbon dioxide and methane or reduced directly to methane using hydrogen as an electron donor (1, 2). Multiple, mechanistically distinct types of C_1_-dependent CoM-methylating enzyme complexes are known in methanogenic archaea.

Activation of methanol or methylamines is mediated by a 3-component system, which produces methyl-CoM via two distinct methyl transfer reactions designated methyltransferase I (MT1) and methyltransferase II (MT2). MT1 enzymes are heterodimeric complexes that transfer the methyl moiety from the substrate to the cob(I)alamin cofactor of their corrinoid-binding subunit, generating a protein-bound methyl-cob(III)alamin intermediate. A distinct MT2 protein then methylates CoM using this protein-bound intermediate as a methyl donor (3–7). The corrinoid-binding subunits of MT1 enzymes comprise a homologous family of proteins related to B12-dependent methionine synthases, whereas the substrate-specific methyltransferase subunits are unrelated to each other (8). In contrast, MT2 methyltransferases comprise a family of homologous proteins with relatively broad substrate specificity (8, 9).

*Methanosarcina barkeri* is known to initiate catabolism of dimethylsulfide (DMS) or methylmercaptopropionate (MMPA) via a 2-component system in which a bifunctional protein homologous to MT2 enzymes, designated MtsA, catalyzes both half-reactions via a ping-pong mechanism (10, 11). In the “MT1” reaction, the corrinoid-binding protein MtsB is methylated by MtsA using a methylsulfide donor. In the “MT2” reaction, MtsA methylates CoM using methyl-MtsB as the C_1_ donor. The chemical similarity of the substrate (DMS or MMPA) and product (methyl-CoM), both of which contain thioether linkages, provides a satisfying explanation for the ability of the MtsA to catalyze both half-reactions (11). This is in stark contrast to activation of methylamines or methanol by 3-component systems, in which the MT1 and MT2 half reactions are chemically distinct.

A third type of activation system is found in *Methanosarcina acetivorans*, where genes encoding a family of proteins (MtsD, MtsF and MtsH) comprised of a corrinoid-binding domain fused to an MT2 domain is required for use of methanethiol (MeSH) and DMS (9, 12). Based on the precedent of the *M. barkeri* MtsA/MtsB system and the chemical similarity of the methyl-thiol substrates, it has been suggested that these proteins catalyze a methylthiol:CoM methyltransferase reaction analogous to the 2-component reaction catalyzed by MtsA/MtsB (9, 12). However, in this case, a single polypeptide carries both the methyltransferase and corrinoid-binding domains. Although purified MtsF (also known as CmtA), is capable of methyl transfer from DMS to the corrinoid cofactor, *in vitro* assays suggest that methyl-tetrahydromethanopterin is a superior substrate. This observation led to the idea that MstF acts to bypass the membrane-bound, ion-motive force dependent methyl-tetrahydromethanopterin:CoM methyltransferase encoded by the *mtrA-H* operon (13). Significantly, the genetic and biochemical observations are not mutually exclusive and it has been suggested that this family of proteins may serve to channel C_1_ units from methylsulfides into both the oxidative and reductive branches (*i.e.* tetrahydromethanopterin and CoM, respectively) of the methylotrophic pathway for methanogenesis (9).

Recently, we showed that growth of *M. acetivorans* using MMPA requires two genes, *mtpC* and *mtpA,* which encode members of the MT1 corrinoid-binding subunit and MT2 methyltransferase families, respectively (9). Based on these genetic data, we predicted that MtpA and MtpC would comprise a two-component methylsulfide:CoM methyltransferase similar to MtsA/MtsB (9). Here, we explicitly test this hypothesis through biochemical characterization of MtpA purified from the native host. Contrary to expectations, our data support the conclusion that MtpA is a substrate-specific MMPA:cob(I)alamin MT1 methyltransferase. Co-purification of the MtsF protein with an MtpA/MtpC complex suggests that it may be involved in MMPA metabolism, perhaps functioning as switch that allows methyl-transfer to both CoM and tetrahydromethanopterin.

## RESULTS

### Purification of MtpA

MtpA was purified from the native host after expression from plasmids encoding an affinity tagged allele of *mtpA.* When the tagged allele was expressed in an *mtpA* deletion mutant (WWM903; Table 1), growth on MMPA was similar to that of the wild-type strain, showing that the tagged MtpA protein is fully functional *in vivo*. When MtpA was affinity purified from this host under strictly anoxic conditions, several additional proteins co-eluted with the tagged protein (Fig. S1). Mass spectrometric analysis identified two of these proteins as MtpC and MtsF. To eliminate the possibility of these co-eluting proteins might interfere with downstream assays, we expressed the tagged protein in a Δ*mtsD,* Δ*mtsF,* Δ*mtsH,* Δ*mtpCAP* host (WWM998, Table 1), which allowed isolation of MtpA that was free of additional proteins based on visual inspection of SDS-PAGE gels (Fig S2).

**Table 1.**
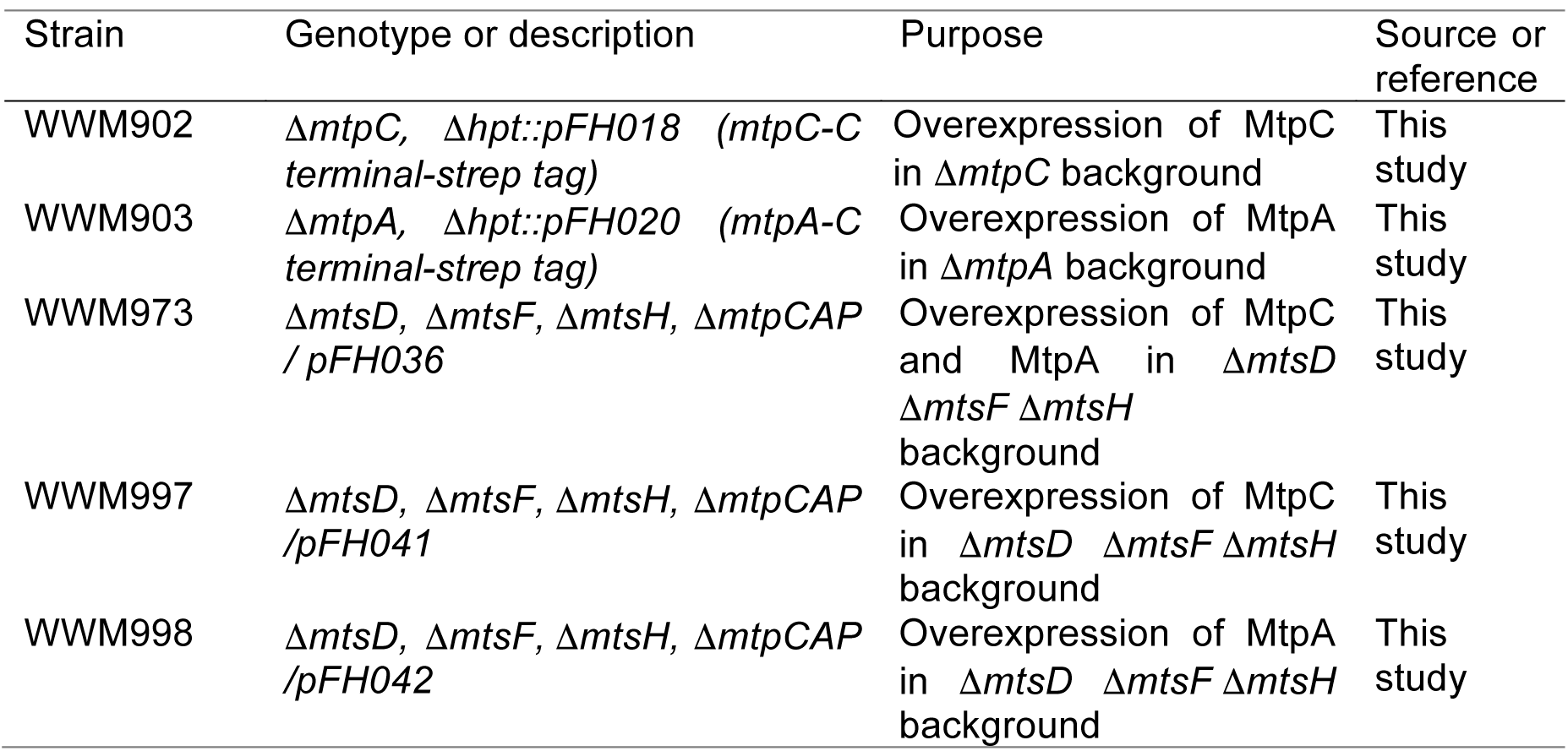
Strains used in this study

| Strain | Genotype or description | Purpose | Source or reference |
| --- | --- | --- | --- |
| WWM902 | $\Delta mtpC$, $\Delta hpt::pFH018$ (*mtpC*-C terminal-strep tag) | Overexpression of MtpC in $\Delta mtpC$ background | This study |
| WWM903 | $\Delta mtpA$, $\Delta hpt::pFH020$ (*mtpA*-C terminal-strep tag) | Overexpression of MtpA in $\Delta mtpA$ background | This study |
| WWM973 | $\Delta mtsD$, $\Delta mtsF$, $\Delta mtsH$, $\Delta mtpCAP$ / *pFH036* | Overexpression of MtpC and MtpA in $\Delta mtsD$ $\Delta mtsF$ $\Delta mtsH$ background | This study |
| WWM997 | $\Delta mtsD$, $\Delta mtsF$, $\Delta mtsH$, $\Delta mtpCAP$ / *pFH041* | Overexpression of MtpC in $\Delta mtsD$ $\Delta mtsF$ $\Delta mtsH$ background | This study |
| WWM998 | $\Delta mtsD$, $\Delta mtsF$, $\Delta mtsH$, $\Delta mtpCAP$ / *pFH042* | Overexpression of MtpA in $\Delta mtsD$ $\Delta mtsF$ $\Delta mtsH$ background | This study |

### MtpA is a specific MMPA:cob(I)alamin methyltransferase

The methyltransferase activity of purified MtpA was assessed using a spectrophotometric assay to monitor the methylation state of corrinoid cofactors and coupled gas chromatography-mass spectrometry (GC-MS) to measure the products and reactants. MtpA catalyzed a rapid methyl transfer reaction using free methylcob(III)alamin as the donor and mercaptopropionate (MPA) as the acceptor (Fig 1). Methyltransferase activity was proportional to protein concentration, while heat-denatured MtpA showed no detectable activity. The stoichiometry of MMPA production and methylcob(III)lamin and MPA consumption was 1:1:1. In the presence of 150 μM methylcob(III)alamin, MtpA displayed an apparent *K*_m_ for MPA of 12.2 ± 1.3 μM, *V*_max_ of 491 ± 12 nmol•min^-1^•mg^-1^ MtpA (*k*_cat_=0.315 s^-1^), *k*_cat_/*K*_m_ =2.6 x 10^4^ M^-1^ s^-1^. Significantly, CoM did not serve as a methyl acceptor under identical conditions (Fig 2). Thus, MtpA does not catalyze an MT2-like reaction.

**Figure 1.**
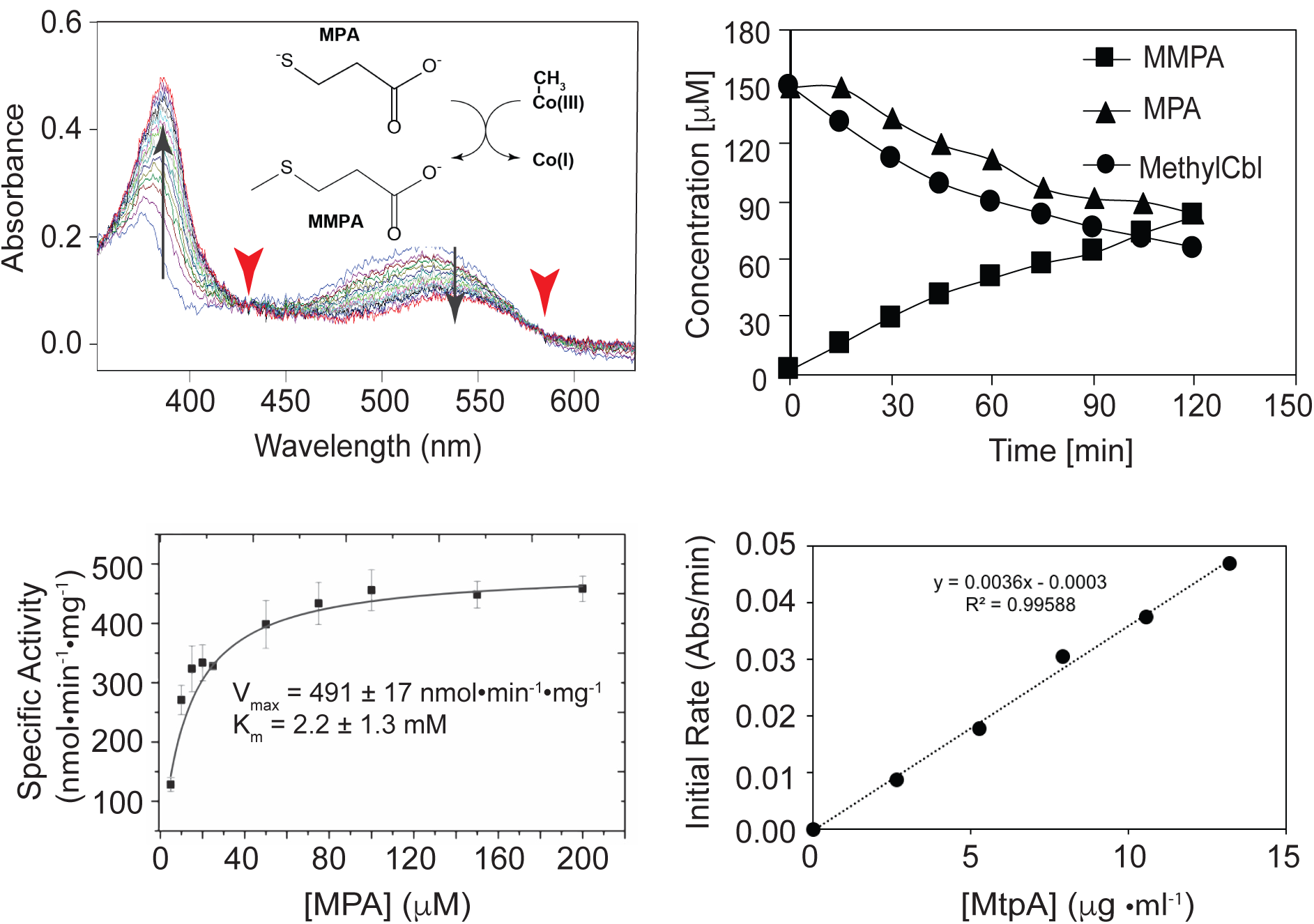
MtpA-catalyzed methylation of MPA using methylcob(III)alamin. *Upper left,* Demethylation of methylcob(III)alamin monitored via UV-visible spectroscopy. The reaction catalyzed is shown in the inset. Spectra were collected at 30 s intervals from 0 min to 8 min. The diagnostic absorbance changes at 388 nm and 540 nm (black arrows) and isosbestic points at 430 nm and 578 nm (red arrowheads) demonstrate the transition from methyl-cob(III)alamin to cob(I)alamin. *Upper right,* Quantification of methylcob(III)alamin, MPA and MMPA during the MtpA-catalyzed reaction. Methylcob(III)alamin was quantified by UV-visible spectroscopy. MPA and MMPA were measured by GC-MS. *Lower left,* Michaelis-Menten kinetics of MtpA. *Lower right,* Methyltransferase reaction is proportional to MtpA concentration. Rates were determined with 150 μM methylcob(III)alamin, 2.7 to 13.5 μg MtpA, 100 mM ZnCl_2_ and 5 mM Ti(III) citrate.

**Figure 2.**
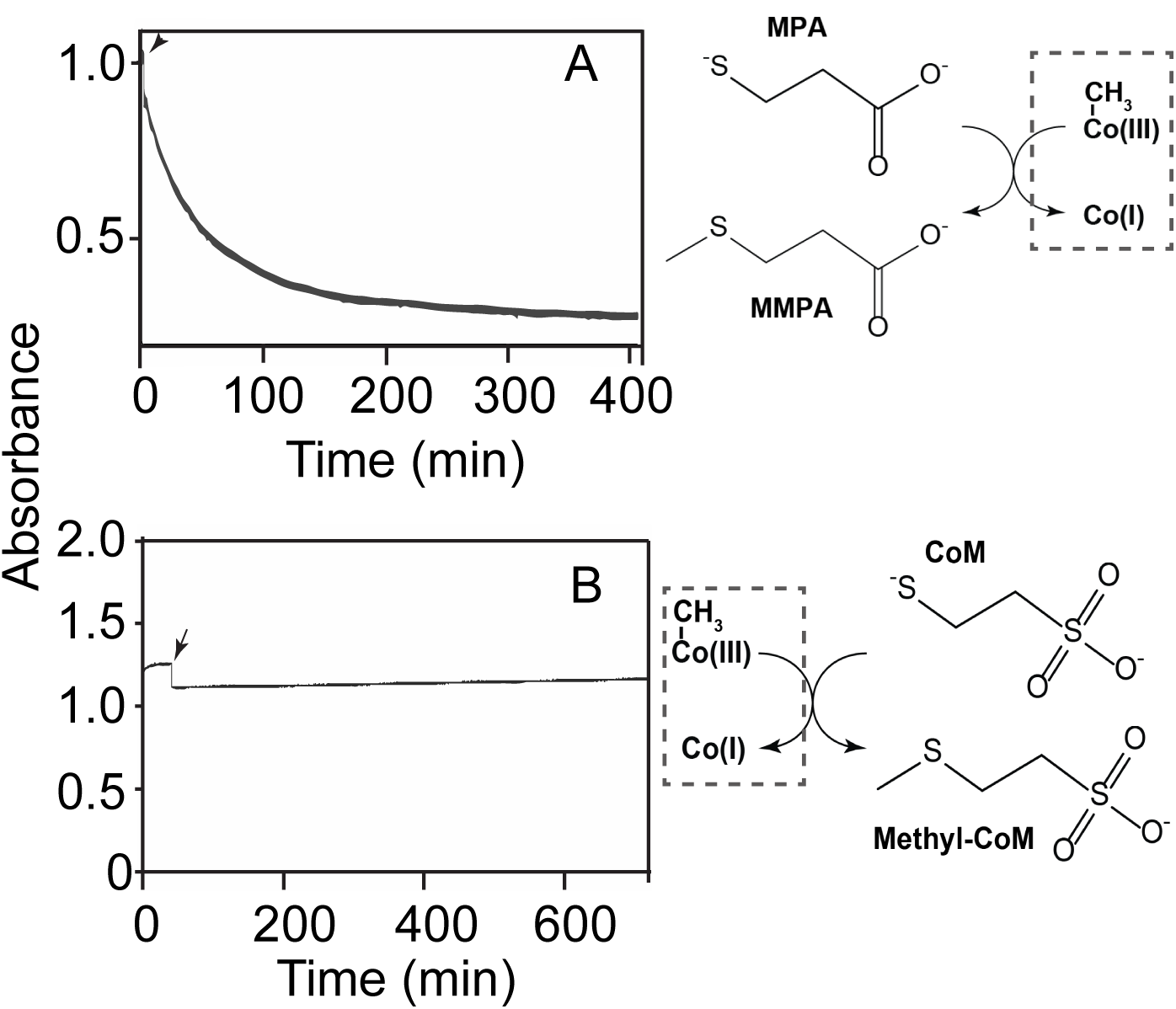
MtpA does not catalyze methylcob(III)alamin:CoM methyl transfer reaction. *Upper panel,* MtpA-catalyzed methylcob(III)alamin:MPA methyltransferase activity monitored by UV-visible spectroscopy at 540 nm. The reaction being monitored is shown to the right of the plot. *Lower panel,* A similar assay attempting to show MtpA-catalyzed methylcob(III)alamin:CoM methyltransferase activity. No activity was observed within the 12-hour assay. Reactions were initiated by addition of substrates as indicated by the black arrows.

Methylation of free cob(I)alamin by MMPA, which is the physiologically relevant direction, was observed only in the presence of very high levels of MMPA (Fig. 3). It should be noted that the requirement for high substrate concentrations is expected due to the unfavorable thermodynamics of this reaction (ΔG^o^’ ≅ 20 kJ/mol, (14)). With 40 mM substrate the initial rate of MMPA:Cob(I)alamin methyl transfer was 32.7 ± 3.8 nmol•min^-1^•mg^-1^(n=3). Methylation of cob(I)alamin was not observed using methanol, dimethylsulfide or dimethylamine as methyl-donors, despite prolonged incubations with high substrate concentrations (Fig. 4).

**Figure 3.**
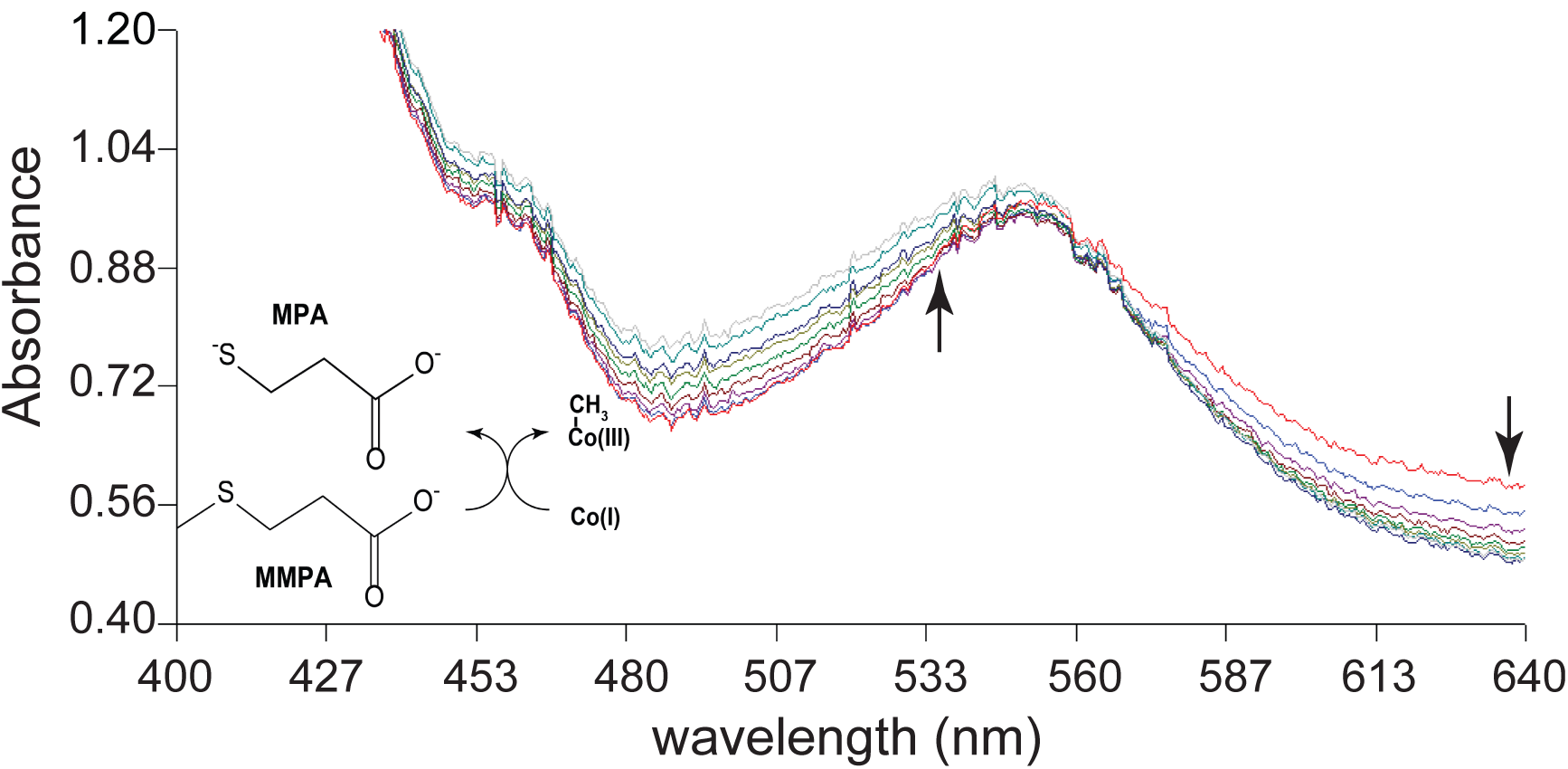
MtpA-catalyzed methylation of Cob(I)alamin methylation using MMPA. Methylation of cob(I)alamin monitored via UV-visible spectroscopy. The reaction catalyzed is shown. Spectra were collected at 10-min intervals from 0 min to 90 min. Black arrows indicate the decrease/increase in absorbance over time.

**Figure 4.**
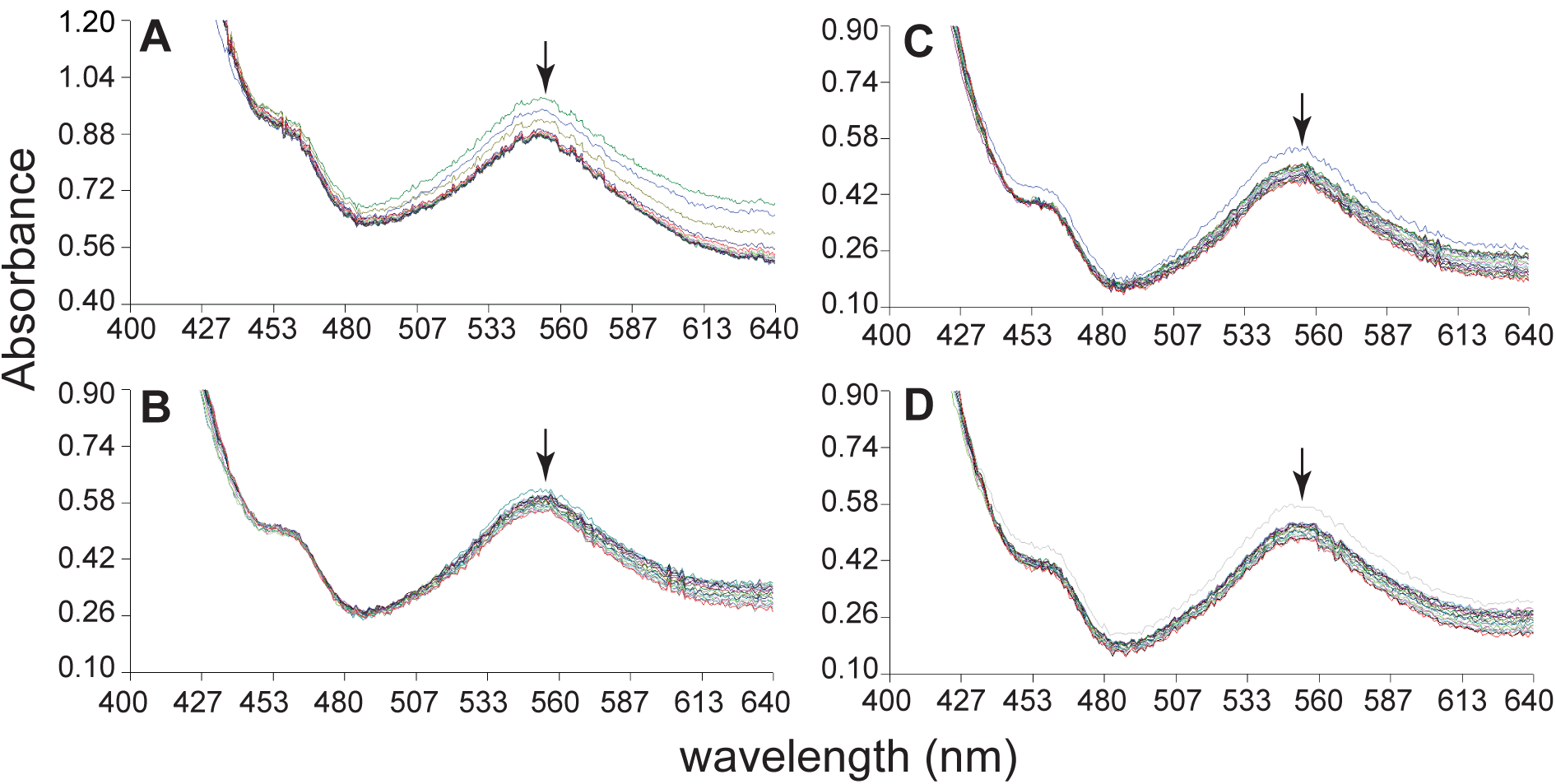
MtpA does not catalyze cob(I)alamin methylation using MeCoM, DMS, MeOH or DMA. Methylation of cob(I)alamin monitored via UV-visible spectroscopy using a variety of potential methyl donors. Formation of methyl-cob(III)alamin would result in an increase in absorbance at 540 nm, as seen in Figure 3, as opposed to the slight loss of absorbance, presumably caused by photobleaching, seen in these assays (indicated by black arrows). *Panel A*. Spectra gathered of cob(I)alamin with 40 mM CH_3_-CoM at 30-min intervals for 10 hours after the start of reaction. *Panel B*. Spectra gathered of cob(I)alamin with 40 mM DMS at 2-min intervals from 0 min to 40 min after the start of reaction. *Panel C*. Spectra gathered of cob(I)alamin with 40 mM MeOH at 2-min intervals from 0 min to 40 min after the start of reaction. *Panel D*. Spectra gathered of cob(I)alamin with 40 mM DMA at 2-min intervals from 0 min to 40 min after the start of reaction.

### Purification and characterization of MtpC and an MtpC/MtpA complex

Affinity-tagged alleles of MtpC were expressed in *M. acetivorans* mutants using plasmids similar to those used for MtpA. A Δ*mtpC* mutant carrying a plasmid expressing C-terminally tagged MtpC (WWM902, Table 1) grew well in media with MMPA as the sole growth substrate. Thus, tagged MtpC is also fully functional *in vivo*. Although the yield of MtpC was very low, we were able to isolate small amounts of affinity-tagged protein from this host. MtpA and MtsF co-eluted with tagged MtpC purified from the Δ*mtpC* mutant (Fig S3). In contrast to our results with MtpA, we were unable to isolate any MtpC after expression in the Δ*mtsD,* Δ*mtsF,* Δ*mtsH,* Δ*mtpCAP* host grown on TMA-medium, suggesting that this protein is unstable in this strain background. However, when tagged versions of both MtpC and MtpA were expressed in the Δ*mtsD,* Δ*mtsF,* Δ*mtsH,* Δ*mtpCAP* host, we were able to purify large amounts of both proteins. The resulting affinity purified protein preparation contained only MtpA and MtpC, which formed a semi-stable complex, as evidenced by an additional band appeared on the native PAGE gel compared to the SDS-PAGE gel (Fig. 5). Taken together, these data suggest that MtpC in unstable in the absence of MtpA.

**Figure 5.**
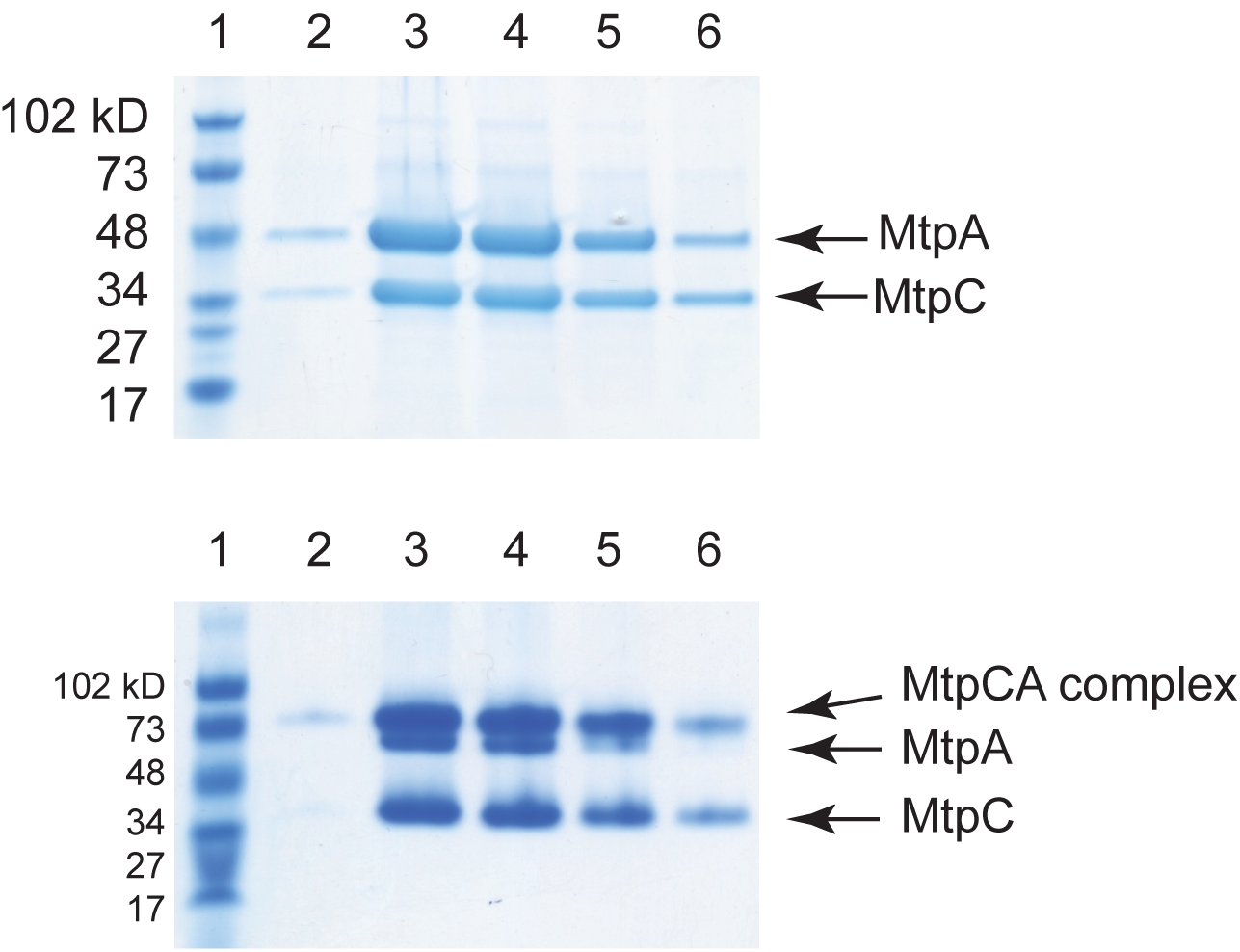
MtpA and MtpC form a complex. Cell-free extract from WWM973 was subjected to affinity purification. The eluted proteins were then separated by electrophoresis and stained with Coomassie Blue. *Upper panel*, a denaturing SDS-PAGE gel (4–20%) was loaded with fractions from different steps of protein purification. Lane 1: protein standard (molecular masses are indicated in kDa to the left); 2-6: elution fractions. *Lower panel*, the same fractions were loaded on a non-denaturing PAGE gel (4-20%). At the pH of running buffer (pH 8.5), both MtpC (pI=4.04) and MtpA (pI=4.52) are negatively charged.

The purified MtpA/MtpC complex exhibited the characteristic spectrum of the corrinoid cofactor in the inactive Co(II) state (Fig. S4) (15). Based on high resolution mass spectrometry, the corrinoid cofactor in MtpC is 5-hydroxybenzymidazolyl cobamide (B_12_HBI, Fig 6), similar to other characterized *Methanosarcina* MT1 enzymes (16–18). We attempted to reactivate the MtpC protein by incubation with strong reducing agents using established protocols that work for some, but not all, corrinoid-dependent methyltransferases. Spectrophotometric analysis suggests that cofactor remained in the Co(II) state throughout a 15 hour incubation. Consistent with this observation, we were unable to demonstrate methyl transfer from MMPA to CoM, nor from Methyl-CoM to MPA, using a sensitive ^1^H nuclear magnetic resonance (NMR) assay and the “reactivated” MtpA/MtpC complex (Fig. S5).

**Figure 6.**
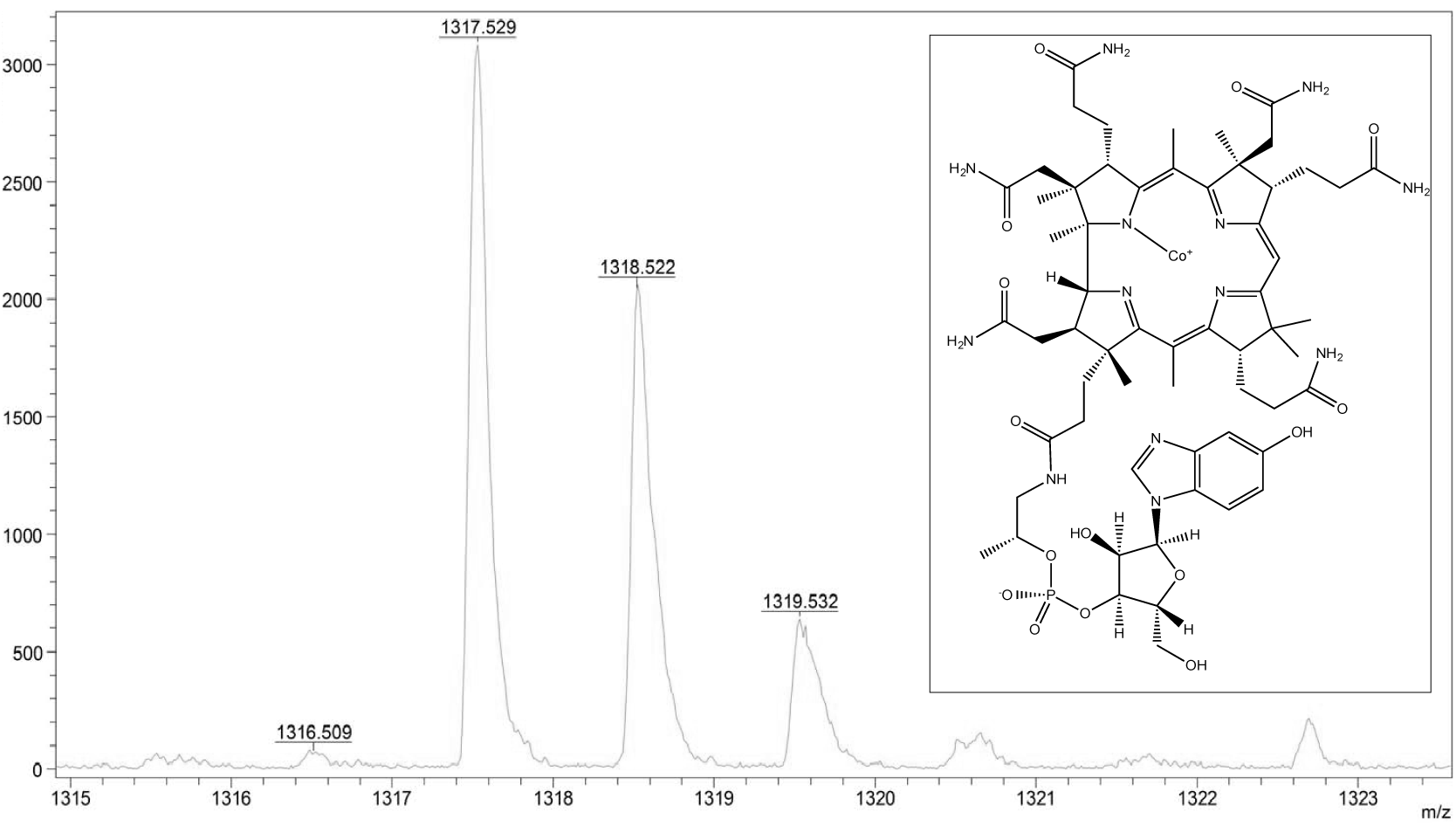
The cofactor of MtpC is 5-hydroxybenzymidazolyl cobamide. High-resolution matrix-assisted laser desorption/ionization mass-spectrometry was used to characterize the corrinoid cofactor purified from the MtpC/MtpA complex. Several masses consistent with 5-hydroxybenzymidazolyl cobamide (shown in the inset) were identified. The predicted exact masses for C_60_H_85_CoN_13_O_15_P (m/z): 1317.535 (100.0%), 1318.539 (64.9%), 1319.542 (20.7%), 1318.532 (4.8%), 1320.545 (3.5%), 1319.539 (3.1%), 1319.536 (2.9%), 1320.543 (2.0%).

## DISCUSSION

MMPA is an abundant molecule that plays an important role in the biogeochemical cycling of both carbon and sulfur. This naturally produced compound is largely derived from degradation of dimethylsulfoniopropionate (DMSP), which is produced in huge quantities by a variety of organisms, most notably marine phytoplankton. Indeed, up to 10% of primary productivity can be directed towards DMSP production in such ecosystems (19). As such, a clear knowledge of the sources and sinks of DMSP is essential for understanding nutrient cycling and ecosystem function on a global scale. Catabolism of DSMP proceeds via two routes. In the first, the molecule is cleaved to release DMS; in the second, it is demethylated to produce MMPA. While the aerobic catabolism of MMPA and DMS has been well studied, our understanding their fate under anoxic conditions has lagged behind (reviewed in (20–22)). Nevertheless, a number of microbes, including methanogenic archaea are known to catabolize both MMPA and DMS. We have been studying one such organism, *M. acetivorans*, in order to better understand the genes and enzymes required for the process.

Bioinformatic and genetic approaches suggested that MMPA catabolism in *M. acetivorans* proceeded via a bifunctional MT1/MT2 MMPA:CoM methyltransferase (9); however, the data presented here clearly show that this is incorrect. Because MtpA catalyzes an MMPA:cob(I)alamin methyltransferase reaction, but not a methylcob(III)alamin:CoM methyltransferase reaction, the MtpA/MtpC complex must be considered an MMPA-specific MT1 enzyme. The high affinity of MtpA for MPA (apparent *K_m_* = 12 μM), coupled with its inability to utilize CoM, supports this conclusion. High substrate affinity is common, but not universal, for substrate-specific MT1 enzymes (*e.g.* the *K_m_* for dimethylamine of MtbB is *ca.* 0.45 mM (23), that of MtgB has for glycine betaine of 1.96 mM (24), while that of MtaB for methanol is *ca.* 40 mM (25). This is in stark contrast to the bifunctional MT1/MT2 activity of the *M. barkeri* MtsA/MtsB complex, which is capable of using a variety of methyl-sulfide substrates, but which has a relatively poor affinity for MMPA (apparent *K_m_* = 10 mM, the K_m_ values for other substrates were not reported) (10, 11). The poor affinity of the bifunctional complex is also very different from the dedicated MT2 enzymes. For example, the MT2 enzymes MtbA and MtaA have much stronger affinities for CoM (*K_m_* 35 μM and or 20 μM, respectively), but very poor affinity for other sulfides like MPA (K_m_ *ca.* 10 mM). Thus, it seems that the bifunctional complex has sacrificed affinity for broad substrate tolerance; however, these data must be interpreted with caution because the K_m_ values for methyl sulfides were determined in the non-physiological, reverse direction due to the fact that the forward reaction is strongly endergonic, with the free energy estimated to be ca. +20 kJ/mol under standard conditions (11). Therefore, the values should be viewed as qualitative, rather than quantitative metrics of substrate preference.

The lack of MT2 activity in purified MtpA implies that an as yet unidentified MT2 is required for MMPA metabolism. The observation that MtsF co-purifies with both MtpA and MtpC raises the possibility that MtsF is this unidentified MT2. This idea is at least partially supported by genetic analysis of *mts* mutants, although the *in vivo* situation is considerably more complex. Thus, while there was no significant phenotype for single or double *mts* mutants, a triple mutant lacking *mtsF* and its two paralogs, *mtsD* and *mtsH* shows a significant defect in both growth rate and yield when grow in MMPA medium (9). These data show that any of the *mts* paralogs suffices for wild-type growth on MMPA. Moreover, the fact that the triple mutant retains some growth on MMPA shows that other MT2 enzymes are capable of fulfilling this function, albeit with lower efficiency than the Mts proteins. In this regard, many MT2 proteins display relaxed substrate specificity. For example, the “methanol-specific” MT2 protein MtaA can serve as the trimethylamine MT2 enzyme both *in vitro* and *in vivo* (26, 27). Similarly, MtsF was previously shown to be involved in metabolism of both DMS (12) and MeSH (9), as well as the transfer of methyl groups from methyl-tetrahydromethanopterin (H_4_MPT) to CoM (13). Because the *M. acetivorans* C2A genome encodes thirteen putative MT2 proteins (8), the comprehensive genetic experiments that will be needed to establish the *in vivo* roles of each MT2 isoenyzme will be challenging.

In the near term, biochemical approaches may be more fruitful for identifying the MT2 protein(s) needed to transfer the methyl group of MMPA to CoM. The goal would be to reconstitute the full MT1/MT2 MMPA:CoM methyltransferase reaction by adding purified MT2 candidates to an active MtpA/MtpC complex. However, this approach will require development of a protocol to convert the inactive Co(II) form of MtaC into the active Co(I) form, which has eluded us to date. Although many corrinoid proteins can be chemically activated using Ti(III), a very low potential electron donor (25, 28), this was not possible for MtpC. Other corrinoid proteins, such as those involved in methylamine catabolism require both Ti(III) and a redox mediator, such as methyl viologen (5–7); however, this treatment also failed to generate the Co(I) form of MtpC. Thus, a different approach will be required to obtain active MtpC. A promising alternative would entail identification of a reductive activation protein, similar to the *Methanosarcina* RamA protein that can be used by to activate the methylamine corrinoid protein (7). We note that a homolog of RamS, denoted RamS, is highly induced by growth on MMPA (9), suggesting a potential path forward for these experiments.

While the data reported here clearly establish the biochemical function of MtpA in methylsulfide metabolism, numerous questions remain. Further, it is becoming clear that the scale and diversity of methylsulfide-dependent methanogenesis has yet to be revealed. Accordingly, a widely distributed and novel order of hydrogen-dependent, methylsulfide metabolizing metabolizing methanogens (Candidatus 'Methanofastidiosa') was only recently discovered (29). These intriguing observations suggest that investigations of methanogenic methylsulfide metabolism will remain fruitful for years to come.

## EXPERIMENTAL PROCEDURES

### Strains, media, and growth conditions

*M. acetivorans* strains were grown in single-cell morphology at 37°C using high-salt (HS) medium containing either 50 mM trimethylamine (TMA) or 20 mM MMPA (30). Cultures used to identify the corrinoid cofactor, were grown is HS medium without added vitamins. Growth on medium solidified with 1.5% agar was as described previously (31). Puromycin was added from sterile, anaerobic stocks at a final concentration of 2 μg/ml for selection of *Methanosarcina* strains carrying the puromycin transacetylase gene (*pac*) (32).

### Chemicals

Ti(III) citrate was prepared anaerobically from TiCl_3_ as described previously (33). Methylthiolpropionate (MMPA) (Tokyo Chemical Industry Co., Japan), and mercaptopropionic acid (MPA) (Sigma, St. Louis, MO) were added from sterile, anaerobic stocks to a final concentration of 20 mM. Methylcob(III)alamin (Sigma, St. Louis, MO) and hydroxocobalamin (Sigma, St. Louis, MO) were prepared as sterile, anaerobic stocks at a final concentration of 1 mM.

### Overexpression of proteins in *M. acetivorans* C2A

Plasmids used for overexpression of MtpC or MtpA were constructed as described in Tables S1 & S2. A C-terminal strep tag was introduced into the MtpC and MtpA coding sequences before cloning into pJK027A to allow affinity purification. The recombinant *mtpC* or *mtpA* plasmids were integrated into the chromosome of corresponding deletion mutants as previously reported (9). To simultaneously overexpress MtpA and MtpC in their native host, a PCR product with both strep-tagged *mtpC* and *mtpA* was cloned into pJK026A, generating pFH035, which was further retrofitted with a pC2A derivative plasmid pAMG40 via the λ-att sites (34), generating the autonomous replicon pFH036.

### Affinity purification of MtpA and MtpC from *M. acetivorans* C2A

*M. acetivorans* strains carrying expression plasmids were grown to late-exponential phase in high-salt medium containing either 20 mM MMPA or 50 mM TMA and 2 μg/mL puromycin. Cells were harvested by centrifugation at 5,000 x g for 15 min at 4°C in sealed anoxic bottles, then brought into a anaerobic chamber (4% H_2_, 96 % N_2;_ Coy Laboratory Products, Grass Lake, MI), where all subsequent manipulations were performed. All buffers used were adjusted pH to 7.2 (unless stated otherwise), made anaerobic by boiling under a N_2_ stream, and brought into the anaerobic chamber in advance. The cells were re-suspended in 10 ml NPD buffer (50 mM NaH_2_PO_4_, 300 mM NaCl, and 2 mM dithiothreitol, pH 8.0) and then sonicated 2 times each for 30 seconds. The cell lysate was centrifuged at 15,000 x g at 4°C for 30 min to remove unbroken cells and debris using a microfuge within the anaerobic chamber. 1ml of Strep-Tactin Superflow Plus (Qiagen) resin was loaded onto Poly-Prep^®^ Chromatography Columns (Bio-Rad, Hercules, CA), and equilibrated with 20 ml anaerobic NPD buffer at 0.5 ml/min. The cell lysate was then loaded onto the column and eluted with 30 ml NPD buffer. The protein was eluted in five fractions with 0.5 ml elution buffer (50 mM NaH_2_PO_4_, 300 mM NaCl, 2 mM dithiothreitol, 2.5 mM desthiobiotin, pH 8.0). The protein concentration in eluted fractions was measured using a Nanodrop 2000 (Thermo Scientific™). The purity of the resultant proteins was judged by visual inspection of samples subjected to by sodium dodecyl sulfate polyacrylamide gel electrophoresis (SDS-PAGE), or native polyacrylamide gel electrophoresis (Native-PAGE), using pre-cast gels (Bio-Rad, Hercules, CA).

### Enzyme assays

All assays were performed inside an anaerobic chamber at room temperature under an atmosphere of 96% N_2_ and 4% H_2_. Data were collected using a fiber optic lead connected to Cary 50 UV-Vis Spectrophotometer. Purified proteins were used immediately after purification and never removed from the anaerobic chamber.

For characterizing the MPA methylation reaction, 1.0 ml reaction mixtures contained 50 mM MOPS/KOH pH 7.2, 100 μM ZnCl_2_, 150 μM methylcob(III)alamin, 5 mM Ti(III) citrate, and 5 μg MtpA. The reaction was started by addition of 150 μM MPA to the mixtures in a cuvette (10-mm light path). The transition from methylcob(III)alamin to cob(I)alamin was monitored by following the decrease in absorbance at 540 nm (ε = 4.4 mM^-1^ • cm ^-1^) (15). The presence of two clear isosbestic points at 430 nm and 578 nm indicated that cob(II)alamin did not accumulate during the reaction. The apparent kinetic parameters of MtpA were investigated by varying the MPA concentration from 7.5 to 150 μM. Results were repeated in triplicate with independently purified batches of MtpA and analyzed using Origin (OriginLab) using a nonlinear Michaelis-Menten parameters.

For methyltransferase assays using various methyl donors and cob(I)alamin as the methyl acceptor, 500 μl reactions were set up in a 10-mm length quartz cuvette (volume=700 μl). The mixtures contained 50 mM MOPS/KOH pH 7.2, 100 μM ZnCl_2_, 0.2 mM hydroxocob(III)alamin, 5 mM Ti(III) citrate, 40 mM methylated substrates (MMPA, DMS, MeOH, DMA or MethylCoM), and 20 μg MtpA. Reactions were initiated by adding respective substrates once the cob(III)alamin was fully reduced to cob(I)alamin, as judged by the UV-visible spectrum. The enzyme-catalyzed transition from cob(I)alamin to methylcob(III)alamin was monitored by following the increase in absorbance at 540 nm. The detection limits of methylcob(III)alamin at 540 nm is 5 μM.

For ^1^H-NMR analysis of MtpC/MtpA activity, 1 ml reactions were set up in a test tube. The mixture contained 50 mM sodium phosphate pH 7.2, 100 μM ZnCl_2_, 5 mM Ti(III) citrate, 0.42 mg MtpC/MtpA, 40 mM substrates (MMPA and CoM, or MethylCoM and MPA). Negative controls used the same amount of heat-killed proteins. The reactions were incubated at room temperature for 16 hours. Samples were passed through 10K Amicon^®^ Ultra filters before ^1^H-NMR analysis.

### Identification of proteins by Mass Spectroscopy

The protein samples were digested by trypsin as follows. Briefly, trypsin (proteomics grade, G-Biosciences) was dissolved in 25 mM ammonium bicarbonate and added at a ratio of 1:20 (trypsin:protein) and processed in a CEM microwave reactor at 55° C for 30 minutes. Digested peptides were lyophilized and separated using a Dionex Ultimate 3000 RSLCnano was connected directly to a Thermo LTQ-Velos-ETD pro mass spectrometer. The column used was an Acclaim 300 C18 nano-column 75µm x 150mm (particle size 3 Å) with an Acclaim Guard column. The flow rate was 300 nl/ml. Typical sample load was 1-2 µg digested peptides and a gradient from 100% A (water + 0.1% formic acid) to 60% B (acetonitrile +0.1% formic acid) was used. Data collection was conducted using the “Big Five” protocol (Thermo, San Jose, CA), MS/MS data were collected using collision induced dissociation (CID). Raw data were collected by Xcalibur (Thermo, San Jose, CA) and processed using an in-house Mascot Distiller and Mascot Server (Cambridge, UK). Resultant peptides generated were searched against NCBI-NR or Uniprot protein databases. Proteins identified that were below the ions score with extensive homology p>0.05 (p is the probability that the observed match is a random event) were discarded. For quantitation, the Exponentially Modified Protein Abundance Index (emPAI) was obtained from Mascot Analysis (35).

### Analysis of cobalamin by Mass Spectroscopy

The corrinoid content of protein samples was determined as described previously (36). Briefly, 0.5 ml protein sample was mixed with 0.7 ml of 95% ethanol and heated to 80°C for 10 min. The mixture was then placed in an ethanol-dry ice bath for 3 min before centrifugation at 10, 000 x g for 10 min. The supernatant was dried under vacuum using a rotary evaporator and re-suspended in 20 μl of double-distilled water before subjecting to analysis using a Bruker UltrafleXtrem MALDI. The measurements were made using the reflection mode and the positive ion was recorded. Samples were prepared by mixing 1 μl of samples and 10 μl of the dihydroxybenzoic acid (DHB) matrix, made from 20 mg of DHB with 1 ml of 50% acetone.

### Determination of metabolites

MMPA and MPA were converted to esters using ethanol before subject to GC-MS analysis. Briefly, 100 µl standard samples (10 μl pentanoic acid was added as an internal control) were mixed with 40 µl pyridine:ethanol (4:1) mixture and 60 µl ethylchloroformate, and incubated at ambient temperature for 45 min inside a fume hood. The bottom phase was extracted into 150 µl dichloromethane after addition of 50 mg NaCl. The samples were dried under N_2_ stream before being dissolved into 100 ul dichloromethane. 5 ul of the resultant samples were injected for GC-MS analysis. Standard samples were prepared on the same day as experimental samples were collected, and were run on the same equipment to generate standard curves. ^1^H-NMR spectra were recorded on an Agilent Technologies DD2 600 MHz spectrometer with D2O as the lock solvent (minimum 10% v/v) at the Carl R. Woese Institute for Genomic Biology.

## Acknowledgements

We thank Dr. Dipti Nayak for constructive suggestions to this project, Dr. Peter Yau from Roy J. Carver Biotechnology Center for protein identification, Dr. Alexander Vladimirovich Ulanov from Metabolomics Center for help with quantification of MMPA and MPA, and Dr. Haijun Yao from Mass Spectrometry Laboratory for technical support with MALDI analysis of cobalamin structure. NMR spectra were recorded on an instrument purchased with support from NIH grant S10 RR028833. This work was supported by National Science Foundation grant MCB-1022462 to W.W.M.

## Author contributions

H.F. performed all protein purification and enzyme assay experiments, analyzed results, and wrote the draft. M.G. performed ^1^H-NMR analysis. W.W.M. supervised overall experimental design, provided critical suggestions, and revised the manuscript. All authors approved the final version of the manuscript.

